# Type III-A CRISPR systems as a versatile gene knockdown technology

**DOI:** 10.1101/2020.09.25.310060

**Authors:** Walter T. Woodside, Nikita Vantsev, Michael P. Terns

## Abstract

CRISPR-Cas systems are functionally diverse prokaryotic anti-viral defense systems, which encompass six distinct types (I-VI) that each encode different effector Cas nucleases with distinct nucleic acid cleavage specificities. By harnessing the unique attributes of the various CRISPR-Cas systems, a range of innovative CRISPR-based DNA and RNA targeting tools and technologies have been developed. Here, we exploit the ability of type III-A CRISPR-Cas systems to carry out RNA-guided and sequence-specific target RNA cleavage for establishment of research tools for post-transcriptional control of gene expression. Type III-A systems from three bacterial species (*L. lactis, S. epidermidis* and *S. thermophilus*) were each expressed on a single plasmid in *E. coli* and the efficiency and specificity of gene knockdown was assessed by Northern blot analysis. We show that engineered type III-A modules can be programmed using tailored CRISPR RNAs to efficiently knock down gene expression of both coding and non-coding RNAs *in vivo*. Moreover, simultaneous degradation of multiple cellular mRNA transcripts can be directed by utilizing a CRISPR array expressing corresponding gene-targeting crRNAs. Our results demonstrate the utility of distinct type III-A modules to serve as effective gene knockdown platforms in heterologous cells. This transcriptome engineering technology has the potential to be further refined and exploited for key applications including gene discovery and gene pathway analyses in additional prokaryotic and perhaps eukaryotic cells and organisms.

## INTRODUCTION

Bacteria and Archaea often harbor CRISPR-Cas systems that provide acquired immunity against viruses and other mobile genetic elements (MGEs) (Hille et al. 2018; Makarova et al. 2019). Upon invasion, a portion of prokaryotic cells integrate short (30-40 base pair) sequences from the MGE genomes into host cell CRISPR (clustered regularly interspaced short palindromic repeat) genomic arrays to provide a heritable record of the captured MGE sequences (Jackson et al. 2017b; McGinn and Marraffini 2019). Immunity requires that the CRISPR arrays are transcribed and the primary CRISPR transcripts processed to generate mature CRISPR (cr)RNAs (Brouns et al. 2008; Carte et al. 2008; Carte et al. 2014). Subsequently, each mature crRNA associates with specific CRISPR-associated (Cas) proteins to form effector crRNPs (crRNA-Cas protein ribonucleoprotein complexes) that mediate crRNA-guided recognition and Cas nuclease-mediated destruction of invasive MGE nucleic acids to prevent further MGE infection (Jackson et al. 2017a; Hille et al. 2018).

CRISPR-Cas systems are diverse and have been categorized into six distinct types (I-VI) that employ different effector Cas nucleases that target recognition and destruction of either foreign DNA (types I, II, V and possibly IV) (Brouns et al. 2008; Garneau et al. 2010; Zetsche et al. 2015; Pinilla-Redondo et al. 2020), RNA (type VI) (Abudayyeh et al. 2016) or both DNA and RNA (type III) (Hale et al. 2009; Staals et al. 2013; Goldberg et al. 2014; Hale et al. 2014; Tamulaitis et al. 2014; Samai et al. 2015; Elmore et al. 2016; Estrella et al. 2016; Kazlauskiene et al. 2016; Han et al. 2017; Tamulaitis et al. 2017). By harnessing the unique attributes of the various CRISPR-Cas systems (such as nucleic acid binding specificity, nuclease activity, etc.), a range of innovative CRISPR-based DNA and RNA targeting tools and technologies have been developed (e.g. for genome editing, control of gene expression, sequence-specific antibiotics, nucleic acid-based viral and pathogen diagnostics) (Bikard et al. 2014; Terns and Terns 2014; Gootenberg et al. 2018; Terns 2018; Pickar-Oliver and Gersbach 2019; Smargon et al. 2020). Tremendous potential remains to exploit our knowledge of various CRISPR systems and components for establishment of innovative research tools with associated transformative biotechnological and biomedical applications.

In this study, we sought to investigate the potential of type III-A CRISPR systems (also referred to as Csm systems (Haft et al. 2005)) to be harnessed as a gene knockdown platform in prokaryotes, akin to how RNA interference (RNAi) machinery has been used in eukaryotes for gene expression downregulation and gene discovery applications (Kim and Rossi 2008; Wilson and Doudna 2013; Setten et al. 2019). Type III-A effector crRNPs are composed of a single crRNA stably associated with five (Csm 1-5) Cas protein subunits (Staals et al. 2014; Tamulaitis et al. 2014; Kazlauskiene et al. 2016; Ichikawa et al. 2017; Liu et al. 2017; Foster et al. 2018; Dorsey et al. 2019; You et al. 2019) (Figure 1A). The mature crRNAs within these complexes are generated through site-specific cleavage of type III-A CRISPR array primary transcripts (within the repeat regions) by the Cas6 endoribonuclease (Carte et al. 2014). The processed crRNAs within the effector crRNPs contain eight nucleotides of repeat sequence at the 5’ end called the 5’ tag (Hale et al. 2009), followed by a ~30-40 nucleotide guide sequence that basepairs with the target RNA protospacer. The 3’ ends of the crRNAs within the complexes can have variable lengths of repeat-derived sequence (Hatoum-Aslan et al. 2013; Tamulaitis et al. 2014; Kazlauskiene et al. 2016). While the repeat-derived, 5’ tag element is critical for function, crRNA species containing or lacking 3’ end repeat sequences are each functional (Hale et al. 2012; Tamulaitis et al. 2014; Foster et al. 2018).

**FIGURE 1.**
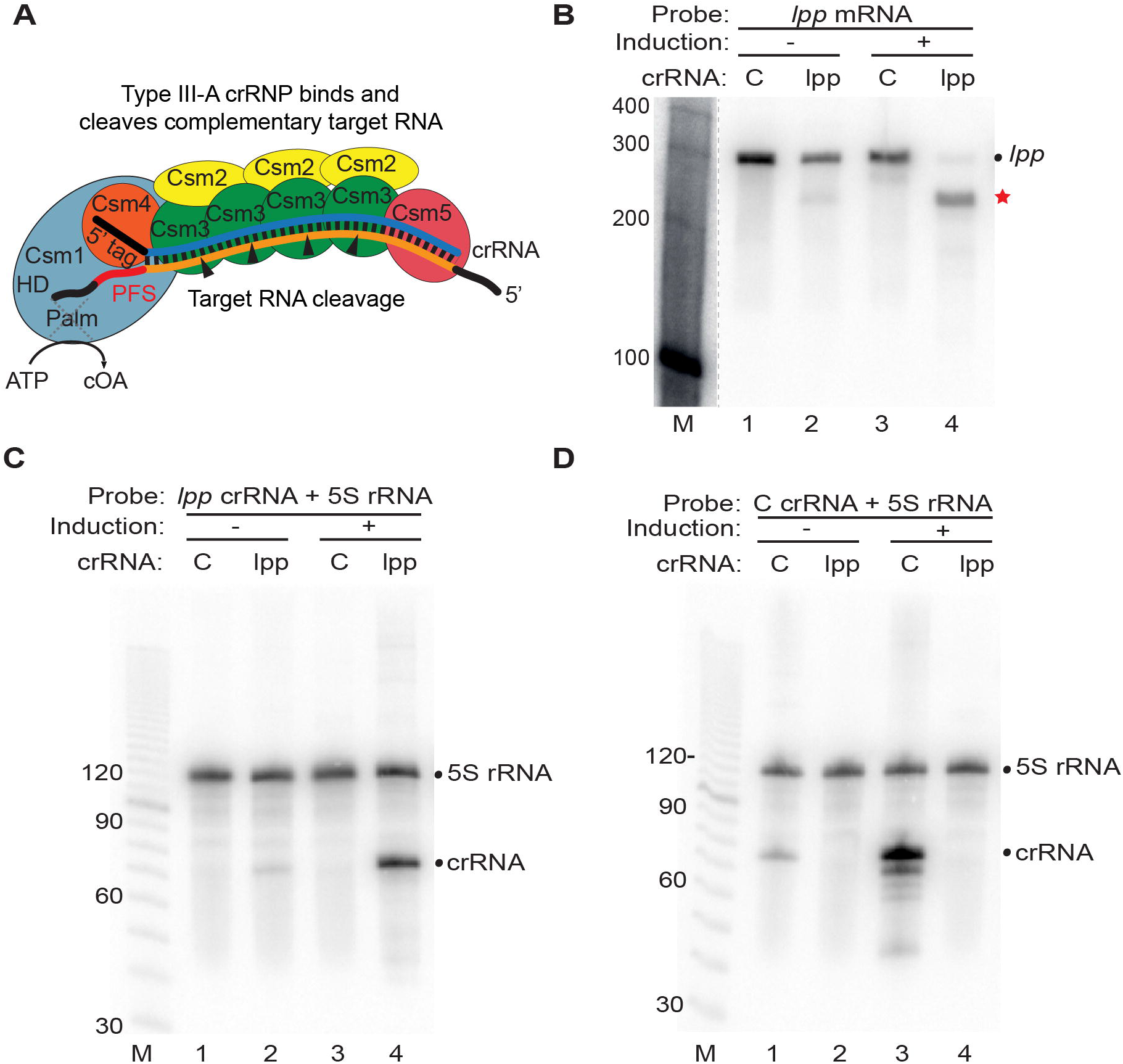
Programmed mRNA cleavage by *L. lactis* type III-A crRNPs expressed in *E. coli.* (A) Diagram of a representative type III-A effector crRNP containing Csm1-5 subunits and a crRNA, in the process of cleaving a bound target RNA. Each Csm3 RNase subunit cuts the target RNA once (cleavages indicated by arrows) within the region of the target RNA (orange) that base-pairs with the crRNA guide element (blue). The position of the target RNA protospacer flanking sequence (PFS) is indicated as are the HD (DNase) and Palm (cyclic oligoadenylate (cOA) producing) motifs of the Csm1 subunit. (B-D) Expression of *L. lactis* type III-A crRNPs containing either a crRNA against the *lpp* mRNA (*lpp*) or negative control crRNA (C) was induced (+) and Northern analysis was performed using probes against the *lpp* mRNA (B), lpp crRNA (C), control crRNA (D) or constitutively expressed 5S rRNA (C and D). The positions of the RNAs are indicated including those of the full-length *lpp* mRNA (dot) and *lpp* mRNA cleavage products (red star). The dotted line in B indicates that intervening lanes were omitted from the blot. The sizes of the molecular weight markers (M) are indicated in each panel.

Type III-A systems can function by both DNA and RNA cleavage mechanisms that are each activated via crRNA-guided recognition of target RNA (Staals et al. 2013; Goldberg et al. 2014; Staals et al. 2014; Tamulaitis et al. 2014; Samai et al. 2015; Jiang et al. 2016; Kazlauskiene et al. 2016; Ichikawa et al. 2017; Kazlauskiene et al. 2017; Liu et al. 2017; Mogila et al. 2019). Target RNA recognition induces conformational changes within the crRNP (Guo et al. 2019; Jia et al. 2019; You et al. 2019) leading to activation of intrinsic DNase and RNase activities as well as triggering production of cyclic oligoadenylate (cOA) from ATP precursors (Kazlauskiene et al. 2017; Niewoehner et al. 2017; Rouillon et al. 2018). In turn, cOA serves as a second messenger that binds to and allosterically activates the RNase activity of a *trans*-acting enzyme called Csm6 (Kazlauskiene et al. 2017; Niewoehner et al. 2017; Foster et al. 2018; Rouillon et al. 2018; Jia et al. 2019), which has been shown to be capable of acting on both target (crRNA-matching) and non-target (cellular) RNAs (Jiang et al. 2016; Rostol and Marraffini 2019). Collectively, these multiple activities lead to a dual ability of type III-A systems to cleave DNA and RNA targets for viral or plasmid clearance or growth arrest or death of infected cells.

The structural organization of type III-A crRNPs and functional roles of the individual Csm1-6 protein subunits have been investigated and three of the subunits have been shown to be catalytic (Rouillon et al. 2013; Hatoum-Aslan et al. 2014; Staals et al. 2014; Tamulaitis et al. 2014; Liu et al. 2017; Jia et al. 2018; Dorsey et al. 2019; Mogila et al. 2019; You et al. 2019). Csm1 (a Cas10 superfamily member (Makarova et al. 2019)) is a large, multiple domaincontaining protein that typically contains two highly conserved functional motifs: the HD motif capable of destroying single-stranded DNA (Elmore et al. 2016; Estrella et al. 2016; Kazlauskiene et al. 2016) and the GGDD motif of one of two Palm domains that can convert ATP into cOA second messenger molecules (Kazlauskiene et al. 2017; Niewoehner et al. 2017; Rouillon et al. 2018; Foster et al. 2020) (Figure 1A). Csm3 is an intrinsic RNA endoribonuclease that cleaves the target RNA in the region defined by crRNA base-pairing at regular six nucleotide intervals (typically 4-5 cuts are made, depending upon the copy number of Cmr3 RNase subunits within the III-A crRNP) (Tamulaitis et al. 2014) (Figure 1A). As noted above, Csm6 is a cOA-activated, trans-acting ribonuclease (Jiang et al. 2016; Niewoehner et al. 2017; Rostol and Marraffini 2019). The DNase (Csm1) and RNase (Csm3 and Csm6) activities of type III-A systems can each contribute to robust immunity against diverse viral and plasmid invaders with notable differences in the dependency on DNase or Csm6-mediated RNase for the various systems investigated (Hatoum-Aslan et al. 2014; Samai et al. 2015; Cao et al. 2016; Foster et al. 2018; Millen et al. 2019; Lin et al. 2020).

Two distinct regions of the target RNA control type III crRNP activities. Extensive complementary binding between the target RNA protospacer region and the crRNA guide region is required to trigger the activities. However, several studies have revealed a key role for the short, 8 nt sequence that flanks the 3’ end of the RNA protospacer, termed the protospacer flanking sequence (PFS), in controlling type III activities (Figure 1A). If the target RNA PFS exhibits perfect or significant complementarity to the 5’ tag element of the crRNA (Pyenson et al. 2017; You et al. 2019); Figure 1A), then target RNA-bound crRNP becomes incapable of triggering DNase activities or cOA production (Marraffini and Sontheimer 2010; Samai et al. 2015; Elmore et al. 2016; Kazlauskiene et al. 2016), but target RNA cleavage is unaffected (Hale et al. 2012; Tamulaitis et al. 2014).

We have transplanted functional III-A systems from three bacterial species (*L. lactis, S. epidermidis* and *S. thermophilus*) by co-expressing the six Csm proteins, Cas6 (for processing of pre-crRNA transcripts into functional crRNAs), and a CRISPR array on a single, arabinose inducible plasmid in *E. coli.* The expressed III-A modules were previously reported to specifically eliminate invading plasmids and anti-plasmid immunity depends on crRNA homology, transcription of the DNA target sequence, cOA signaling and RNase activity of Csm6, rather than on the DNase activity of Csm1 (Ichikawa et al. 2017; Foster et al. 2018). Here, we have specifically exploited the site-specific (Csm3-mediated) RNA cleavage activity of the three distinct type III-A crRNPs to efficiently knock down gene expression of both coding and non-coding RNAs *in vivo.* We demonstrate that the III-A modules can be programmed to recognize one or more *E. coli* cellular target RNAs by addition of appropriate crRNA coding sequences to the module. Our findings demonstrate the potential of heterologous type III-A systems as tools for RNA interference and pave the way for expanded utility as functional genomic tools in novel cells and organisms.

## RESULTS

### *Programming L. lactis* III-A crRNPs for selective cleavage of target mRNAs *in vivo*

To determine if type III-A crRNPs can be harnessed as a gene expression knockdown platform, we introduced a plasmid into *E. coli* cells that results in arabinose-inducible expression of type III-A crRNPs (Ichikawa et al. 2017; Foster et al. 2018) that were programmed with different crRNA sequences rationally designed to recognize specific *E. coli* target mRNAs (Figure 1). The *E. coli* host strain (BL21-AI) lacks endogenous CRISPR-Cas systems. Northern blot analyses, using probes against the target mRNAs as well as internal control 5S rRNA, were carried out to assess the efficiency and specificity of directed cleavage of endogenous *E. coli* mRNAs.

One of two distinct approaches were employed in this study to ensure that sequence-dependent target RNA cleavage was specifically mediated by the backbone RNase subunits, Csm3, and not through triggering the non-specific RNase activity of trans-acting Csm6, which requires cOA-signaling by the crRNPs (Figure 1A). Initially, this goal was accomplished by searching target RNAs for the presence of 3’ PFS sequences with significant homology to the 8 nt 5’ crRNA tag. Pairing between the crRNA tag and PFS of the target RNA results in Csm3-directed cleavage of crRNA matching target RNAs but prevents the complex from producing cOA second messenger required for activating non-specific Csm6 RNase activity (Kazlauskiene et al. 2016; Pyenson et al. 2017; You et al. 2019). As a more versatile strategy, we subsequently employed type III-A crRNPs that have a mutation in the conserved Csm1 Palm motif (GGDD to GGAA) shown to be incapable of producing cOA/Csm6 activation regardless of the sequence nature of the PFS in the desired target RNA (Kazlauskiene et al. 2017; Niewoehner et al. 2017; Rouillon et al. 2018; Foster et al. 2020). Some type III-A systems employ a non-specific DNase activity of the Csm1 HD domain (Jiang et al. 2016; Kazlauskiene et al. 2016; Park et al. 2017), but we previously reported that the systems investigated here do not rely on this activity for immunity against plasmid DNA and therefore we did not employ Csm1 HD mutants (Foster et al. 2018). Mutation of the Palm or HD motif of Csm1 does not influence the ability of the type III-A crRNPs to cleave target RNA (Figure S1).

For the initial test, we assessed the RNA targeting capacity of heterologously expressed, *L. lactis* type III-A crRNPs having a crRNA against the *lpp* mRNA (the *lpp* gene of *E. coli* is nonessential for viability and encodes a major lipoprotein (Baba et al. 2006)). The *lpp* mRNA target sequence was selected with perfect complementarity to the crRNA guide and to have a 3’ PFS with extensive base-pairing potential with the 5’ crRNA tag (PFS bases −1 to −5 and −8 were complementary to positions +1-5 and +8 of the 5’ crRNA tag) to ensure that only the Csm3-mediated specific target RNA cleavage (and not Csm6 non-specific RNase activity) was triggered. As a specificity control, the same *L. lactis* crRNPs were programmed with a control crRNA (C) that does not contain significant complementarity to any *E. coli* RNA. In the presence of the crRNA against the *lpp* mRNA but not the non-cognate control crRNA (Figure 1B, compare lanes 3 and 4), we observed a significant reduction in the steady-state levels of the *lpp* mRNA that was accompanied by the appearance of a *lpp* mRNA breakdown product of the expected size for cleavage in the region recognized by the crRNA (Figure 1B, red star in lane 4). Furthermore, crRNA-dependent cleavage of the target *lpp* mRNA was dependent upon arabinose induction of expression of the type III crRNP (Figure 1B, compare lanes 2 and 4). As expected, the steady state levels of 5S rRNA remained constant in the analyzed cells (Figures 1C and D). These same samples were also probed to confirm the expression of each engineered crRNA. The size of the main crRNA species matches that expected for the product following Cas6 cleavage of primary transcripts (71 nt) and low levels of presumably 3’ trimmed crRNAs was also observed (*lpp* crRNA is shown in Figure 1C, lanes 2 and 4 and non-cognate control crRNA is shown in Figure 1D, lanes 1 and 3). Mutational analysis of Csm3 (D30A) confirmed that the RNase activity observed by the crRNPs is mediated by the Csm3 backbone subunit cutting within the target RNA protospacer region, as expected (Figure S2). The results show that heterologously expressed *L. lactis* type III-A crRNPs can be programmed to efficiently and selectively cleave a desired target mRNA *in vivo.*

### *S. thermophilus* and *S. epidermidis* type III-A systems can also be programmed to target mRNA destruction

To determine if type III-A crRNPs from other bacterial species were also capable of carrying out efficient and specific mRNA knockdown in *E. coli,* we programmed the *S. thermophilus* and *S. epidermidis* type III systems to target the same *lpp* endogenous mRNA transcript (Figure 2). Similar to the *L. lactis* system (Figure 1), our results revealed that expression of III-A crRNPs from both *S. thermophilus* (Figure 2A-C) and *S. epidermidis* (Figure 2D) resulted in specific and efficient knockdown of the *lpp* mRNA and the accumulation of mRNA cleavage products of the expected sizes for crRNA-directed cleavage. While *L. lactis* and *S. epidermidis* share the same 5’ crRNA tag sequence (5’-ACGAGAAC-3’), the *S. thermophilus* 5’ tag differs at positions 4 and 5; 5’-ACGGAAAC-3’). For this reason, we used a Csm1 Palm mutant (GGDD motif mutated to GGAA) for the source of *S. thermophilus* crRNPs to enable testing of the same target RNA region and to ensure that Csm6 non-specific RNase activity was not triggered due to decreased tag-PFS complementarity. Thus, three distinct bacterial sources of type III-A CRISPR-Cas systems proved to be functional for programmable RNA targeting *in vivo.*

**FIGURE 2.**
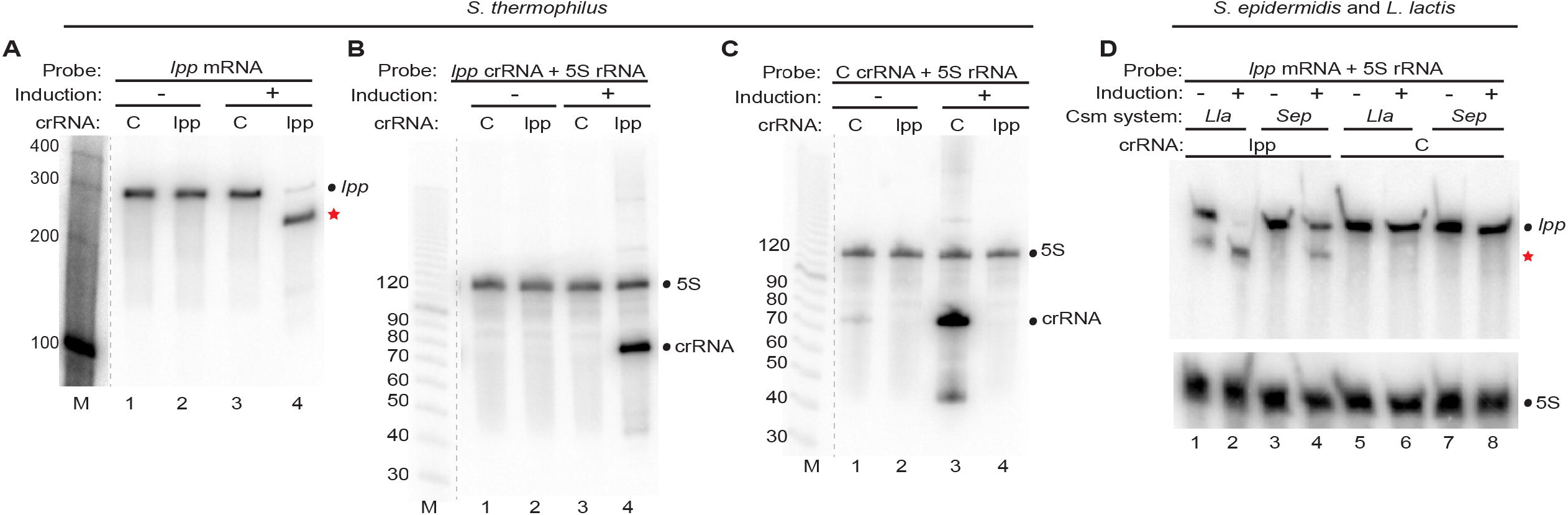
Targeted mRNA destruction by *S. thermophilus* and *S. epidermidis* type III-A crRNPs. (A-C) Expression of *S. thermophilus* type III-A crRNPs containing either a crRNA against the *lpp* mRNA (*lpp*) or negative control crRNA (C) was induced (+) and Northern analysis was performed using probes against the *lpp* mRNA (A), lpp crRNA (B), control crRNA (C) or 5S rRNA (B and C). (D) Expression of the *S. epidermidis (Sep*) and *L. lactis (Lla*) type III-A crRNPs containing either a crRNA against the *lpp* mRNA or control crRNA was induced (+) and Northern analysis was performed using probes against the *lpp* mRNA and 5S rRNA. The positions of the RNAs are indicated including those of the full-length *lpp* mRNA (dot) and *lpp* mRNA cleavage products (red star). (A-C) The dotted lines indicate intervening lanes were omitted from the blot. The sizes of the molecular weight markers (M) are indicated in each panel.

### Type III-A crRNPs are capable of acting at different sites along the length of a target mRNA

Next, we investigated whether the position of the target site significantly impacts the efficacy of target mRNA cleavage by testing the ability of the *L. lactis* III-A crRNPs (with Csm1 GGDD motif mutation) to target cleavage at different sites along the *lpp* mRNA transcript. Specifically, we individually tested the effects of five distinct crRNAs that span different regions of the *lpp* mRNA and in each case, Northern analyses was performed with probes specific for either the 5’ or 3’ terminal regions of the *lpp* mRNA (Figure 3A). Each of the tested crRNAs led to a major reduction in the steady state levels of full-length *lpp* transcript relative to the control crRNA, when probing for either the 5’ or 3’ ends of the mRNA following induction of crRNP formation (Figures 3B and C). Moreover, 5’ and 3’ expected cleavage products were observed with each tested crRNA (Figures 3B and C). Following crRNA-guided target RNA cleavage, the 3’ mRNA fragments exhibited generally higher steady state levels than 5’ mRNA fragments, despite near full cleavage of the full-length transcripts for each of the tested crRNAs. The results indicate that type III crRNPs are capable of targeting mRNA cleavage at multiple sites along the length of transcripts.

**FIGURE 3.**
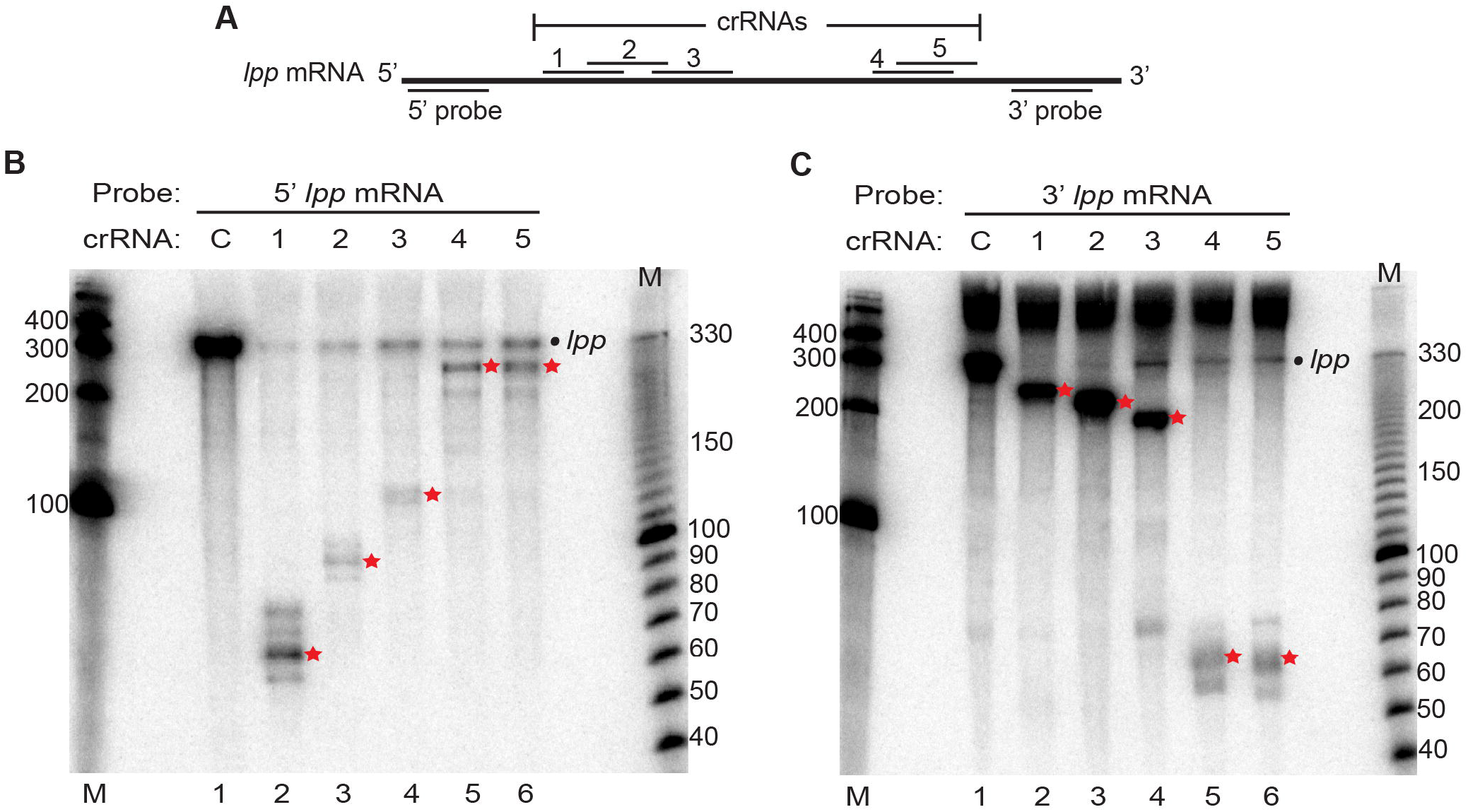
Type III-A crRNPs are effective at targeting cleavage at distinct sites along the length of a mRNA. (A) Diagram of the *lpp* mRNA showing the relative positions of each tested crRNA and Northern probes employed. (B-C) Expression of the *L. lactis* type III-A crRNPs containing different crRNAs (1-5) was induced and Northern analysis was performed using probes against the 5’ (B) and 3’ (C) termini of the *lpp* mRNA. The positions of the full-length *lpp* mRNA (dot) and cleavage products (red star) are indicated in each panel. The sizes of the molecular weight markers (M) are indicated in each panel.

### Type III-A crRNPs can be programmed to cleave multiple distinct RNA targets simultaneously

We next determined if the *L. lactis* III-A crRNPs could be programmed to target destruction of mRNAs in addition to the *lpp* transcript and addressed if more than one mRNA could be targeted in the cell concurrently. We found that *L. lactis* crRNPs efficiently targeted the destruction of two additional tested non-essential *E. coli* mRNAs (Baba et al. 2006) encoding either the cold-shock protein E (*cspE*) or the outer membrane protein F (*ompF*), when individually programmed with single crRNAs against each of these particular transcripts (Figure 4B, lane 3 and 4C, lane 4; *lpp* mRNA cleavage shown in Figure 4A, lane 2). Next, we attempted targeting *lpp*, *cspE,* and *ompF* mRNAs simultaneously upon expressing each of the three distinct crRNAs from a single CRISPR array. A similar reduction in full-length transcript levels and occurrence of expected cleavage products was observed in the strain expressing all three crRNAs compared to individual strains expressing just one crRNA against a single mRNA target (Figure 4; compare lanes 2 and 5 in A, lanes 3 and 5 in B, and 4 and 5 in C). These results demonstrate the capacity of type III-A crRNPs to carry out multiplexed and simultaneous knockdown of at least three distinct mRNAs.

**FIGURE 4.**
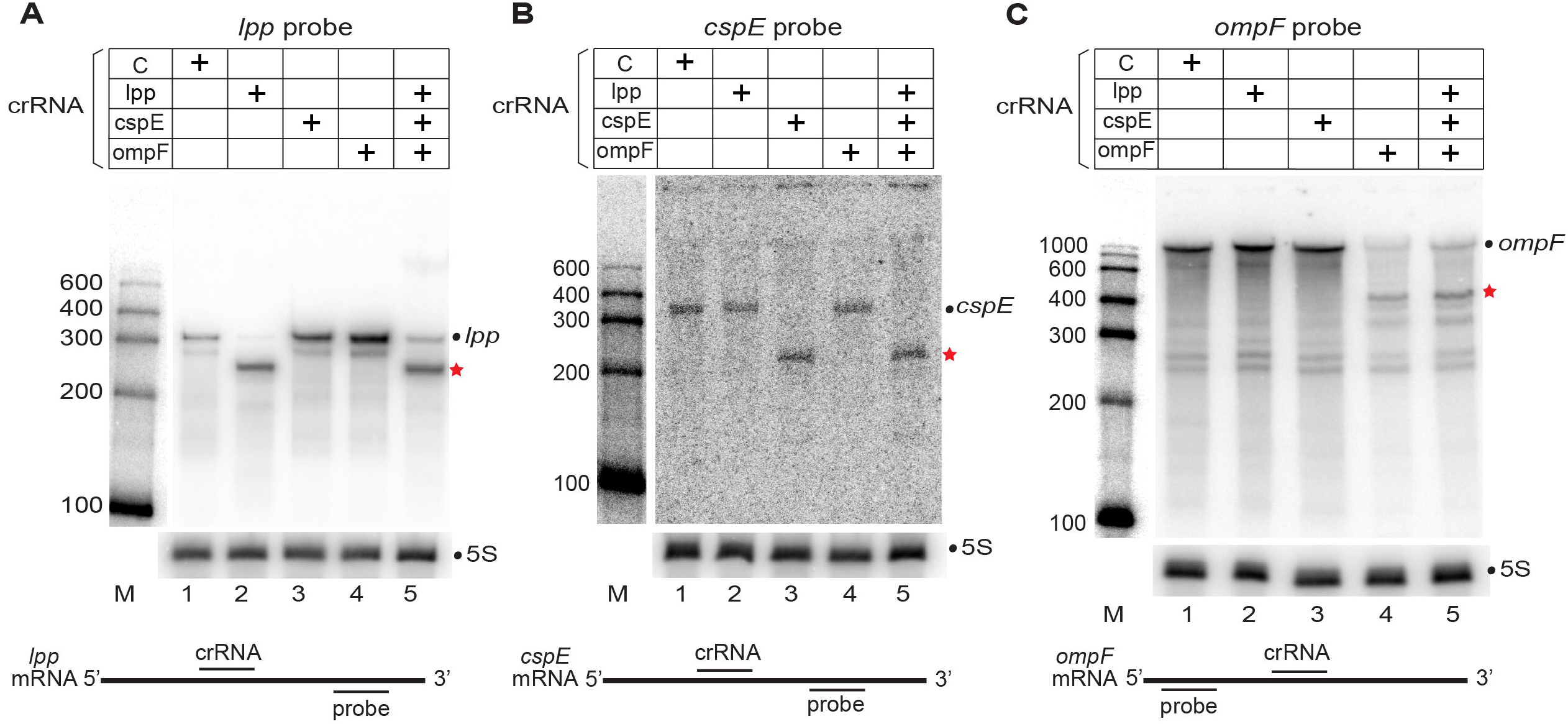
Type III-A crRNPs can effectively target multiple mRNA transcripts simultaneously. (A-C) Expression of the *L. lactis* type III-A crRNPs with single or multiple crRNAs were induced and Northern analysis was performed with probes against *lpp* mRNA (A), *cspE* mRNA (B), *ompF* mRNA (C) or 5S rRNA (A-C). The positions of full-length mRNAs (dot) and expected cleavage products (red star) are indicated in each panel. The sizes of the molecular weight markers (M) are indicated in each panel. The relative positions of the crRNA tested and Northern probe used for each mRNA is shown below each panel.

### Directed cleavage of a non-coding RNA target

To potentially broaden the application beyond mRNA targeting, we next tested whether *L. lactis* III-A crRNPs (with Csm1 Palm mutation) could effectively cleave a non-coding RNA target. For this purpose, we chose to target destruction of the *rnpB* RNA/ribozyme which is the enzymatic component of RNase P and catalyzes 5’ end cleavage of precursor tRNAs (Esakova and Krasilnikov 2010). In *E. coli,* the *rnpB* RNA is required for the endonucleolytic separation of the *valV-valW* pre-tRNA bicistronic transcript (Mohanty and Kushner 2007; Mohanty et al. 2020). We individually tested a panel of seven different crRNAs that spanned the length of *rnpB* RNA (Figure 5 B) for their ability to direct *L. lactis* III-A crRNPs to destroy the RNase P RNA (Figure 5C) and lead to decreased processing of the bicistronic pre-tRNA transcript (Figure 5E).

**FIGURE 5.**
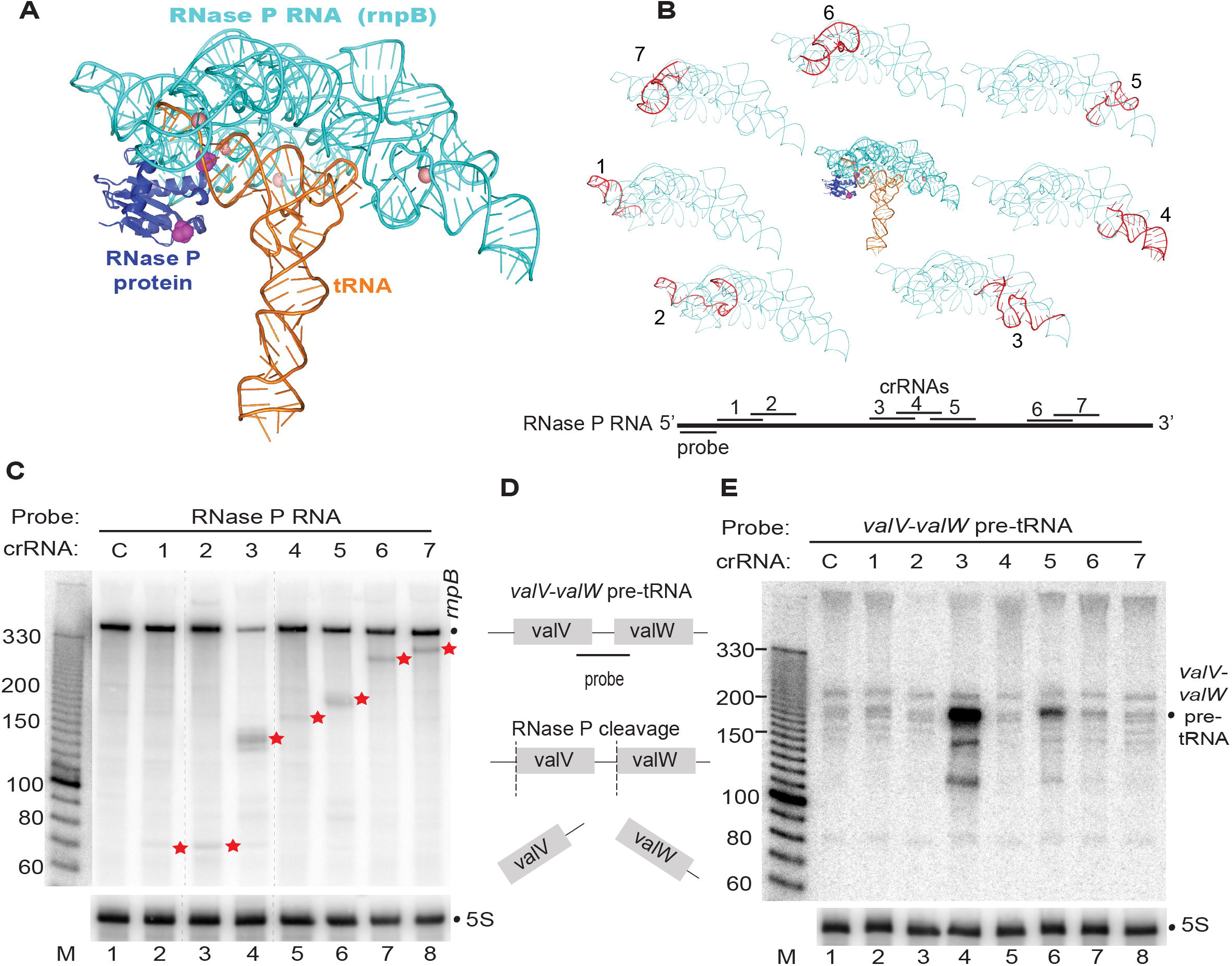
Targeting the non-coding RNA component of RNAse P. (A) Structure of the *T. maritima* RNAse P RNA (*rnpB* RNA in cyan) in complex with the RnpA protein (purple) and substrate tRNA (orange) (PDB 3Q1R). (B) The relative positions of each tested crRNA (1-7) is indicated both on the lower diagram as well as superimposed (red) onto the *rnpB* RNA structure (cyan). (C, E) Expression of the *L. lactis* type III-A crRNPs containing a crRNA targeting *rnpB* (1-7) were induced and Northern analysis was performed with probes against the *rnpB* RNA (C), *valV-valW* pre-tRNA (E) or 5S rRNA (C and E). The positions of the *rnpB* RNA full-length (dot) and cleavage products (red stars) and sizes of molecular weight markers (M) are indicated. (D) Schematic of the *valV-valW* pre-tRNA processing by RNase P and binding location of probe used to selectively recognize the uprocessed pre-tRNA transcript (E).

Northern analysis revealed that there was a high variability in the ability of the selected crRNAs to significantly lower the steady-state levels of *rnpB* RNA (Figure 5C). All seven tested crRNAs exhibited the ability to cleave the *rnpB* RNA as evidenced by the accumulation of expected size cleavage products, relative to the control crRNA (Figure 5C, compare lanes 2-8 with lane 1). However, only one of the seven tested crRNAs resulted in a significant reduction in the steadystate levels of the *rnpB* target RNA (Figure 5C, lane 4) and a corresponding accumulation of unprocessed *valV-valW* bicistronic transcript (Figure 5E, lane 4). Above background levels of bicistronic pre-tRNA accumulation was also observed for a second tested crRNA (Figure 5E, lane 6). The *E. coli* RNase P RNA exhibits a high degree of sequence conservation with the RNase P RNA of *T. maritima,* which has a solved three-dimensional X-ray structure of the folded *rnpB* RNA in complex with tRNA substrate and associated RNase P protein (Reiter et al. 2010) (Figure 5A). We note that the two most effective crRNAs (3 and 5) each are predicted to bind to regions of folded *rnpB* ribozyme that are not interacting with pre-tRNA or RNase P protein (Figure 5B). These results indicate the feasibility of using type III-A crRNPs for knocking down expression of non-coding RNAs but indicate that the folded structure and/or protein binding of a target RNA may negatively impact the effectiveness of target RNA recognition by type III-A crRNPs.

## DISCUSSION

Gene knockdown technologies such as RNA interference (RNAi), play an important role in gene function discovery (especially for essential genes that when knocked out cause lethality) and therapeutic applications (Kim and Rossi 2008; Wilson and Doudna 2013; Setten et al. 2019). The RNAi machinery required for small interfering RNA (siRNA) or short hairpin RNA (shRNA) mediated gene knockdown is present in many eukaryotes but absent from prokaryotes. In this study, we sought to harness the type III-A CRISPR system as a post-transcriptional gene knockdown platform that functions in prokaryotic cells. Our proof-of-principle studies demonstrate that type III-A crRNPs from three distinct sources (*L. lactis, S. epidermidis,* and *S. thermophilus*) and conveniently expressed from an ‘all-in-one’ single plasmid, can be readily programmed with rationally-designed crRNAs to cleave both mRNA and non-coding RNA *in vivo* in a heterologous prokaryotic (*E. coli*) host cell. Moreover, we show that more than one target mRNA can be efficiently cleaved simultaneously when multiple crRNAs are concurrently expressed, paving the way for important applications such as cellular pathway discovery and manipulation. The type III-A gene knockdown technology established here has future potential as an important functional genomics tool for investigating gene function and cellular pathways in a wide range of other prokaryotic and perhaps eukaryotic cells.

### Programmable knockdown of diverse RNA transcripts

Our findings show that type III-A systems provide a versatile platform for controlling the levels of endogenous transcripts *in vivo* at a post-transcriptional level. We found that each of the crRNAs that we designed and tested were highly effective at reducing the steady-state levels of all tested mRNAs when assayed individually (Figures 1-4) or in a multiplexed fashion (Figure 4). Furthermore, crRNAs targeting various locations along the length of a mRNA were comparably effective (Figure 3) suggesting no major bias of particular mRNA regions for susceptibility to interaction and cleavage by III-A crRNPs. Because the guide elements of the type III-A crRNAs that we tested are naturally relatively long (35-37 nt), this likely contributes to the observed effectiveness of targeting specific transcripts *in vivo.* When multiple crRNAs were expressed simultaneously, a high degree of sequence-specificity was observed in targeting the efficient destruction of each target RNA (Figure 4). However, future transcriptomic experiments are required to more comprehensively determine if any of the tested crRNAs result in any off-target effects on the levels of other cellular transcripts.

In contrast to the efficient targeted destruction of specific mRNAs when several crRNAs were tested, the non-coding RNA component of RNase P (*rnpB*) proved to be more recalcitrant to type III-A crRNP-mediated RNA cleavage (Figure 5). Given that each of the panel of seven crRNAs tested did lead to some detectable level of directed cleavage, this indicates that crRNPs were able to recognize and cleave at least a small fraction of the ribozyme. Of note, the two crRNAs found to be most effective at reducing ribozyme levels and blocking pre-tRNA processing function, mapped to regions of the RNA that are predicted to be solvent exposed in the predicted three-dimensional structure and not located in regions known to be engaged in known RNA-protein or RNA-tRNA interactions (Figure 5). It remains to be determined if noncoding RNAs will generally be more refractory to III-A-mediated RNA knockdown or if this particularly highly folded and highly interactive RNA target is an exceptional case. A prudent general strategy for targeting either a specific mRNA or non-coding RNA of interest would be to pursue the multiplexing route. Simultaneous expression of multiple crRNAs against different regions of the desired target RNA molecule is expected to increase the probability of efficient cleavage and gene knockdown.

### Target RNA cleavage products accumulate *in vivo*

The observation that target RNA cleavage products of the expected sizes relative to the site of crRNA interaction were readily detectable provides strong evidence that the destruction was directed by type III-A crRNPs. However, this phenotype was surprising given *a priori* expectations that cleavage of the phosphodiester bonds within RNA polynucleotide chains normally leads to rapid degradation of the cleavage fragments and given the short half-lives (typically 3-5 mins) of mRNAs in *E. coli* in general (Bernstein et al. 2002). It is unclear why RNA fragments located both 5’ and 3’ to the site of cleavage persist and are readily observable under steady state conditions. Type III crRNP cleavage results in RNA products having 5’ hydroxyl and 2’,3’-cyclic phosphate moieties rather than more typical 5’ phosphate and 3’ hydroxyl ends created by the action of other RNases (Hale et al. 2009; Zhang et al. 2016). These particular RNA chemical end groups might impede further RNA turnover by *E. coli* exoribonucleases. Alternatively, the unexpected stability of the 5’ and 3’ degradation fragments might result from the III-A crRNP sterically protecting these ends from being recognized and destroyed by cellular ribonucleases. However, strong *in vitro* evidence revealed that type III crRNPs normally rapidly dissociate from target RNAs following cleavage making this possibility less likely (Estrella et al. 2016; Rouillon et al. 2018).

### Comparison with other RNA targeting CRISPR systems

Most types of CRISPR systems (types I (Cas3), II (Cas9), V (Cas12)) act through crRNA guided Cas nucleases that destroy DNA targets (Hille et al. 2018; Makarova et al. 2019). In contrast, type III (Csm3 and Csm4) and type VI (Cas13) CRISPR systems naturally recognize and target sequence-specific cleavage of RNA substrates. Thus, there has been a recent push to develop CRISPR-based systems as much needed, RNA targeting research tools with novel applications (Terns 2018; Smargon et al. 2020).

Previous work showed that related type III-B (also known as Cmr (Haft et al. 2005)) systems could be programmed with engineered crRNAs to guide destruction of target RNAs in hyperthermophilic archaea including *Pyrococcus* and *Sulfolobus* species that naturally contain the type III-B systems (Hale et al. 2012; Zebec et al. 2014; Liu et al. 2018). In these systems, a CRISPR mini-array plasmid supplies the engineered crRNAs and the endogenous III-B crRNPs are relied upon for the knockdown activity, which requires extreme temperature (optimal growth temperatures of these extremophilic organisms are 70-100 °C). The type III-A systems that we have established potentially have greater utility for use in a broad range of important prokaryotes given that the entire system (crRNP and crRNAs) has been encoded in one plasmid. Importantly, the activity of the type III-A crRNPs described here work at mesophilic (e.g. 37 °C) temperatures compatible with the optimal growth temperatures of many prokaryotic model organisms currently being investigated.

As a tool for RNA knockdown in prokaryotes, type III-A systems have a distinct advantage over the type IV systems which suffer a significant drawback in that the relevant Cas13 RNase is known to cleave both the target RNA as well as “bystander” RNAs (i.e. cellular RNAs in a sequence-nonspecific fashion, when activated by crRNA-target RNA interaction (Gootenberg et al. 2017; Smargon et al. 2017; Konermann et al. 2018; Meeske et al. 2019)). Interestingly, this “collateral” RNA destruction observed with most Cas13 ribonucleases in bacteria was unexpectedly suppressed when certain Cas13 species were tested in human and plant cells and resulted in efficient and specific knockdown of target mRNAs with fewer off-target effects than is observed with siRNA/shRNA technologies (Abudayyeh et al. 2017; Konermann et al. 2018; Wessels et al. 2020). Furthermore, while the type III-affiliated Csm6 (III-A) and Csx1 (type III-B) ribonucleases are also known to induce collateral RNA destruction (Gootenberg et al. 2018; Rostol and Marraffini 2019), these proteins work *in trans* and are not required for the sequence-specific RNA cleavage mediated by the type III crRNPs (Hale et al. 2012; Hale et al. 2014; Tamulaitis et al. 2014). To circumvent collateral RNA cleavage and direct only sequencespecific target RNA destruction, in this work we prevented Csm6 RNase activity through mutating the Csm1 Palm motif essential for cOA generation and Csm6 RNase activation. However, it should also be possible to achieve this same effect by deleting the *csm6* genes from the expression plasmids and we show that deletion of csm6 gene or expression of an RNase-defective Csm6 variant (HEPN mutant) do not impact target RNA cleavage by type III-A crRNPs (Figure S2).

### Future promising applications for type III-A systems

Further studies are required to more deeply explore the potential of the type III-A crRNPs as effective posttranscriptional gene knockdown platforms as well as for development of additional applications. Moreover, it will be important to determine if these CRISPR research tools are capable of functioning outside of *E. coli* and in a range of both prokaryotic and eukaryotic cells and organisms. The ability to assemble functional crRNPs through expression of all required components on just a single expression plasmid, makes this a facile system for further refinement and testing these possibilities. Recent success with ectopically expressing similarly complex, type I crRNPs consisting of up to six Cas proteins and a CRISPR RNA in human cells capable of function in genome editing/transcriptional control (Cameron et al. 2019; Pickar-Oliver et al. 2019) offers hope that type III-A systems too will be able to be employed in eukaryotic cells for important *in vivo* applications.

The ease by which both type III-A and III-B crRNPs can be expressed and purified as functional complexes (Hale et al. 2012; Staals et al. 2013; Staals et al. 2014; Tamulaitis et al. 2014; Osawa et al. 2015; Elmore et al. 2016; Estrella et al. 2016; Kazlauskiene et al. 2016; Foster et al. 2018; Dorsey et al. 2019; You et al. 2019) offers additional opportunities for delivering programmed, pre-assembled crRNPs directly into cells. Of note, *S. thermophilus* type III-A crRNPs expressed and purified from *E. coli* have recently been shown to efficiently and specifically knockdown maternal mRNAs when microinjected into early zebra fish embryos (Fricke et al. 2020). Moreover, as has been employed with type VI, Cas13-based systems, there is the potential to employ ribonuclease defective type III-A crRNP variants (through inactivating point mutations to create an RNase-defective or dCsm3 subunit (Figure S2; (Tamulaitis et al. 2014; Samai et al. 2015)) as well as to fuse various effector domains from other proteins to expand the functionalities beyond RNA destruction to potentially influence target RNA splicing, base editing, translation, degradation, and to track the intracellular localization of the transcripts using fluorescent-based microscopy of GFP-fusion systems (Terns 2018; Smargon et al. 2020). Purified type III-A crRNPs also offer potential to expand proven CRISPR-based powerful molecular diagnostic tools (so far employed with Cas13 and Cas12) capable of detecting viral and bacterial pathogens and cancer mutations from patient fluids (Gootenberg et al. 2017; Gootenberg et al. 2018b; Kellner et al. 2019). In summary, the work described here provides an important step in the direction of harnessing the potential of type III-A systems as versatile RNA-targeting CRISPR-based research tools with important future applications.

## MATERIALS AND METHODS

### Plasmid Construction

pCsm plasmids for expressing *L. lactis, S. epidermidis,* or *S. thermophilus* type III-A crRNPs were described previously (Ichikawa et al. 2017; Foster et al. 2018) with the new crRNAs designed to be complementary to a particular target RNA region. For each crRNA guide sequence (Table S1), a pair of complementary oligonucleotides, 35 bp for *L. lactis* and *S. epidermidis,* 39 bp for *S. thermophilus,* and 37 bp for the negative control (Table S2), was designed with 4 nt 5’ overhangs that match 5’ overhangs of the pCsm vector left by linearization with *BbsI* (NEB). 10 pmoles of each oligonucleotide set were annealed in 1X CutSmart buffer (NEB) and 0.1 pmole of the annealed products were ligated with 50 ng of the linearized pCsm vector with T4 DNA ligase (NEB). The multispacer array was constructed by amplification of individual spacer arrays with oligonucleotides that were appended with BsaI (NEB) restriction sites such that PCR fragments could be combined in a Golden Gate Assembly (Table S2). The ligation and assembly reactions were used to transform chemically competent TOP10 *E. coli* cells (Thermo Fisher). Plasmids were purified (ZymoPURE mini-prep, Zymo Research) and verified by DNA sequencing before transformation of chemically competent BL21-AI *E. coli* cells (Invitrogen). The BL21-AI *E. coli* expression strain has a T7 RNA polymerase under the control of an arabinose inducible promoter.

Splicing overlap extension PCR was used to generate the *L. lactis* Csm1 Palm (D576A, D577A) and HD (and H13A, D14A) motif mutations as well as the *S. thermophilus* Csm1 Palm (D575A, D576A) motif. The gel purified DNA fragments were digested with PspXI and NdeI (*L. lactis*) or BamHI and NdeI (*S. thermophilus*) before ligation into linearized pCsm vector. *L. lactis* Csm3 D30A, ΔCsm6, and Csm6 H360A mutations were transferred from previously generated constructs (Foster et al. 2018) by digestion and ligation. The oligonucleotide sequences used to generate the mutant constructs are provided in Supplementary Table S3 and all mutations were verified by DNA sequencing.

### Type III-A crRNP expression

Single *E. coli* colonies were grown at 37°C shaking in Miller’s lysogeny broth (Invitrogen) until reaching an OD_600_ of 0.1 when they were induced with a final concentration of 10 mM arabinose to express Csm1-6, Cas6, and crRNA. After 120 minutes of induction, 1-1.5 ml of each culture was centrifuged, the supernatant was aspirated, and the cell pellets were flash frozen in a dry ice and ethanol bath and stored at −80C until RNA extraction.

### Northern analysis

Total RNA was prepared using the RNAsnap™ protocol (Stead et al. 2012). RNA used in Figure 5, was further purified by phenol chloroform isoamyl alcohol extraction (pH 4.5) and ethanol precipitation. The RNA samples were quantified using a Qubit fluorimeter 2.0 (Thermo Fisher) and equal amounts of RNA (5 −10 ug) were heat denatured for 5 minutes at 95 °C immediately before electrophoresis on 6-8% polyacrylamide, 8M urea gels in TBE buffer (89 mM Tris base, 89 mM Boric acid, 2mM EDTA, pH 8.0) at 400 volts. 5’-radiolabeled, molecular weight markers were RiboRuler Low Range RNA Ladder (Thermo Scientific) or 10 bp DNA Ladder (Invitrogen). The RNA was transferred by electro-blotting with a semidry transfer apparatus (Bio-Rad® Trans-Blot SD) to positively charged nylon membrane (Nytran™ SPC, Whatman). ULTRAhyb buffer (Invitrogen) was used for hybridization with oligonucleotides 5’-end labeled with 6,000 Ci/mmol γ-^32^P ATP (Perkin Elmer) using T4 polynucleotide kinase (NEB). 1 million counts per minute of radiolabeled probe was added for each mL of hybridization buffer. Hybridization was performed at 42°C for 12-16 hours. Membranes were washed of unbound probe with a pre-warmed (42°C) 2X saline-sodium citrate (SSC) buffer and detected by phosphor imaging (Storm™ 840, GE Healthcare).

## ACKNOWLEDGEMENTS

We are grateful to Sidney Kushner and Bijoy Mohanty for expert technical advice, helpful discussions, and probe sequence recommendations. We thank members of the Terns laboratory for helpful discussions and Rebecca Terns for her early mentorship role. Ryan Catchpole is gratefully acknowledged for critical reading of the manuscript. This work was supported by the National Institutes of Health [R35GM118160] to M.P.T

## SUPPLEMENTARY DATA

Supplementary data are available.

