## Supplemental files for "Type III-A CRISPR systems as a versatile gene knockdown technology"

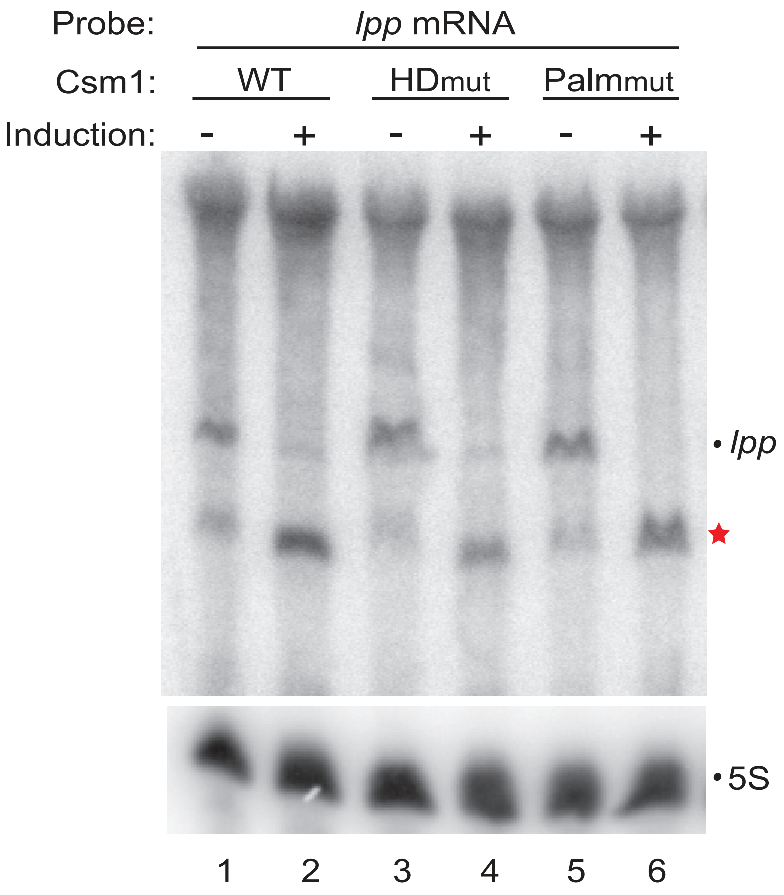

**Supplemental Figure S1.** Mutagenesis of conserved HD and Palm motifs of Csm1 does not impact target RNA cleavage. Expression of *L. lactis* type III-A crRNPs containing either wildtype (WT) or mutant (HD or Palm mutants as indicated) Csm1 and crRNA targeting *lpp* mRNA was induced with arabinose (+) and Northern analysis was performed with probes against the *lpp* mRNA and 5S rRNA. The positions of both the full-length (dot) and cleavage product (red star) of the *lpp* mRNA are indicated.

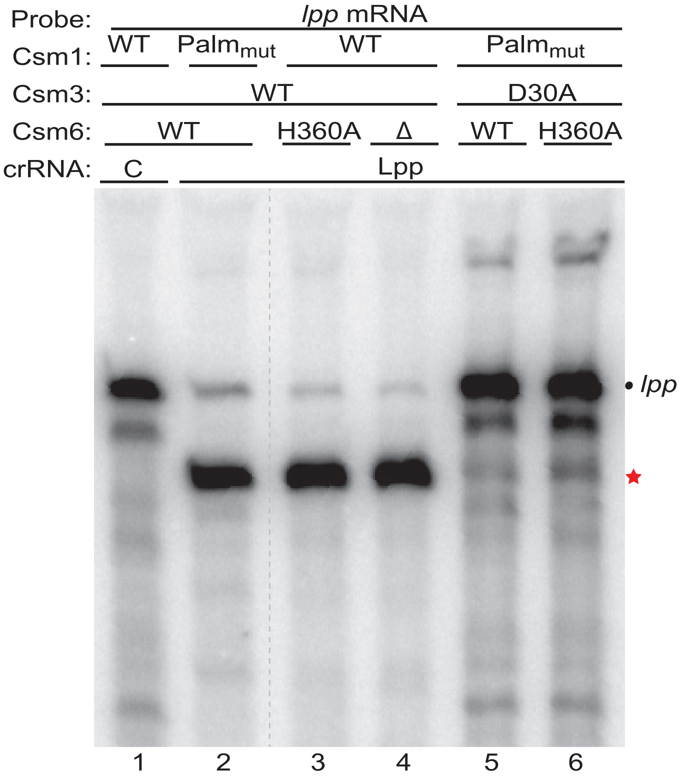

**Supplemental Figure S2.** Mutagenesis of Csm3 (D30A) inhibits target RNA cleavage. The expression of *L. lactis* type III-A crRNPs containing the indicated Csm1, 3 and 6 variants with either the *lpp* crRNA or negative control crRNA (C) was induced with arabinose and Northern analysis was performed with a probe against the *lpp* mRNA. The positions of both the full-length (dot) and cleavage product (red star) of the *lpp* mRNA are indicated. The dotted line indicates the removal of an intervening lane.

**Table S1. Guide sequences for each tested target RNA**

| Target | System | crRNA | Guide sequence 5'-3' |
|---|---|---|---|
| lpp | *L. lactis* | lpp 1 | CCAGGAUUACCGCGCCCAGUACCAGUUUAGUAGCU |
| lpp | *L. lactis* | lpp 2 | AGCAACCUGCCAGCAGAGUAGAACCCAGGAUUACC |
| lpp | *L. lactis* | lpp 3 | ACAGCUGAUCGAUUUUAGCGUUGCUGGAGCAACCU |
| lpp | *L. lactis* | lpp 4 | GCGGUAUUUAGUAGCCAUGUUGUCCAGACGCUGGU |
| lpp | *L. lactis* | lpp 5 | CUUGCGGUAUUUAGUAGCCAUGUUGUCCAGACGCU |
| lpp | *S. epidermidis* | lpp 1 | CCAGGAUUACCGCGCCCAGUACCAGUUUAGUAGCU |
| lpp | *S. thermophilus* | lpp 1 | CCAGGAUUACCGCGCCCAGUACCAGUUUAGUAGCUUUCA |
| cspE | *L. lactis* | cspE 1 | GGACUCAUUAAACCACUUAACGUUACCUUUAAUCU |
| ompF | *L. lactis* | ompF 1 | CAACGUCAGCGUAUUUAAGACCCGCGAAUGCCAGA |
| rnpB | *L. lactis* | rnpB 1 | UAUGGAGCCCGGACUUUCCUCCCCUCCGCCCGUCU |
| rnpB | *L. lactis* | rnpB 2 | CCAUCGGCGGUUUGCUCUCUGUUGCACUGGUCGUG |
| rnpB | *L. lactis* | rnpB 3 | CCGGGUUCAGUACGGGCCGUACCUUAUGAACCCCU |
| rnpB | *L. lactis* | rnpB 4 | CCCGUCUCCCCCGAAGAGGACGACGACGAAGCGGC |
| rnpB | *L. lactis* | rnpB 5 | CCUGAUCCCGCUUGCGCGGGCCAUCGGCGGUUUGC |
| rnpB | *L. lactis* | rnpB 6 | UGCGCUCUUACCGCACCCUUUCACCCUUACCUGAU |
| rnpB | *L. lactis* | rnpB 7 | CUCACUGGCUCAAGCAGCCUACCCGGGUUCAGUAC |

**Table S2. Oligonucleotides used to make crRNA guides**

| pCsm | crRNA | ID | Oligonucleotide sequence 5'-3' |
|---|---|---|---|
| *L. lactis* | lpp 1 | 3025 | GAACCCAGGATTACCGCGCCCAGTACCAGTTTAGTAGCT |
| | | 3026 | ATTTAGCTACTAAACTGGTACTGGGCGCGGTAATCCTGG |
| *S. epidermidis* | lpp 1 | 3025 | GAACCCAGGATTACCGCGCCCAGTACCAGTTTAGTAGCT |
| | | 3142 | CGATAGCTACTAAACTGGTACTGGGCGCGGTAATCCTGG |
| *S. thermophilus* | lpp 1 | 4670 | AAACCCAGGATTACCGCGCCCAGTACCAGTTTAGTAGCTTTCA |
| | | 4671 | TATCTGAAAGCTACTAAACTGGTACTGGGCGCGGTAATCCTGG |
| *L. lactis* | C | 3205 | GAACTTGTAGTATGCGGTCCTTGCGGCTGAGAGCACTTCAG |
| | | 3206 | ATTTCTGAAGTGCTCTCAGCCGCAAGGACCGCATACTACAA |
| *S. thermophilus* | C | 4388 | AAACTTGTAGTATGCGGTCCTTGCGGCTGAGAGCACTTCAG |
| | | 4389 | TATCCTGAAGTGCTCTCAGCCGCAAGGACCGCATACTACAA |
| *L. lactis* | lpp 2 | 3146 | GAACAGCAACCTGCCAGCAGAGTAGAACCCAGGATTACC |
| | | 3147 | ATTTGGTAATCCTGGGTTCTACTCTGCTGGCAGGTTGCT |
| *L. lactis* | lpp 3 | 3098 | GAACACAGCTGATCGATTTTAGCGTTGCTGGAGCAACCT |
| | | 3099 | ATTTAGGTTGCTCCAGCAACGCTAAAATCGATCAGCTGT |
| *L. lactis* | lpp 4 | 4686 | GAACGCGGTATTTAGTAGCCATGTTGTCCAGACGCTGGT |
| | | 4687 | ATTTACCAGCGTCTGGACAACATGGCTACTAAATACCGC |
| *L. lactis* | lpp 5 | 4684 | GAACCTTGCGGTATTTAGTAGCCATGTTGTCCAGACGCT |
| | | 4685 | ATTTAGCGTCTGGACAACATGGCTACTAAATACCGCAAG |
| *L. lactis* | cspE 1 | 3649 | GAACGGACTCATTAAACCACTTAACGTTACCTTTAATCT |
| | | 3650 | ATTTAGATTAAAGGTAACGTTAAGTGGTTTAATGAGTCC |
| *L. lactis* | ompF 1 | 3152 | GAACCAACGTCAGCGTATTTAAGACCCGCGAATGCCAGA |
| | | 3153 | ATTTTCTGGCATTCGCGGGTCTTAAATACGCTGACGTTG |
| *L. lactis* | rnpB 1 | 4009 | GAACTATGGAGCCCGGACTTTCCTCCCCTCCGCCCGTCT |
| | | 4010 | ATTTAGACGGGCGGAGGGGAGGAAAGTCCGGGCTCCATA |
| *L. lactis* | rnpB 2 | 4011 | GAACCCATCGGCGGTTTGCTCTCTGTTGCACTGGTCGTG |
| | | 4012 | ATTTCACGACCAGTGCAACAGAGAGCAAACCGCCGATGG |
| *L. lactis* | rnpB 3 | 4013 | GAACCCGGGTTCAGTACGGGCCGTACCTTATGAACCCCT |
| | | 4014 | ATTTAGGGGTTCATAAGGTACGGCCCGTACTGAACCCGG |
| *L. lactis* | rnpB 4 | 5212 | GAACCCCGTCTCCCCCGAAGAGGACGACGACGAAGCGGC |
| | | 5213 | ATTTGCCGCTTCGTCGTCGTCCTCTTCGGGGGAGACGGG |
| *L. lactis* | rnpB 5 | 5214 | GAACCCTGATCCCGCTTGCGCGGGCCATCGGCGGTTTGC |
| | | 5215 | ATTTGCAAACCGCCGATGGCCCGCGCAAGCGGGATCAGG |
| *L. lactis* | rnpB 6 | 5216 | GAACTGCGCTCTTACCGCACCCTTTCACCCTTACCTGAT |
| | | 5217 | ATTTATCAGGTAAGGGTGAAAGGGTGCGGTAAGAGCGCA |
| *L. lactis* | rnpB 7 | 5218 | GAACCTCACTGGCTCAAGCAGCCTACCCGGGTTCAGTAC |
| | | 5219 | ATTTGTACTGAACCCGGGTAGGCTGCTTGAGCCAGTGAG |

Paired oligonucleotides were annealed and ligated into linearized pCsm to program guide RNAs

| | | | |
|---|---|---|---|
| Multispacer array | lpp 1 | 3721 | CAGCAAGGTCTCAGAACCCAGGATTACCGC |
| | | 3722 | GATGCTGGTCTCAGGAGCGGTTGTATTTAGCTAC |
| | cspE 1 | 3723 | CAGCAAGGTCTCGCTCCTCGATAAAAGGGGACGAGAACGGAC |
| | | 3724 | GATGCTGGTCTCCCCTTTTATCGAGGAGCGGTTGTATTTAGAT |
| | ompF 1 | 3725 | CAGCAAGGTCTCAAAGGGGACGAGAACCAAC |
| | | 3726 | GATGCTGGTCTCGCGGTTGTATTTTCTGGCATT |

Paired oligonucleotides were used to amplify fragments from single spacer array plasmids, which were then combined in a Golden Gate Assembly into pCsm.

**Table S3. Oligonucleotides used to make mutations**

| Mutation | ID | Oligonucleotide sequence 5'-3' |
|---|---|---|
| *L. lactis* Csm1 H13A D14A D576A D577A | 3197 | AAGCAACCTCGAGGCTGTGGTCTA |
| | 3186 | GTGCCACGATAGATAATTTTGCCAATAGCAGCCAGCAGGCTACCACAAACC |
| | 3187 | ATTGGCAAAATTATCTATCGTGGCAC |
| | 3188 | CGCCACCGGCATAAATAAC |
| | 3189 | GTTATTTATGCCGGTGGCGCCGCCCTGTTTATGATTGGTGCATGGC |
| | 3198 | AAAAGCCATATGTTAGCGTTCACG |
| *S. thermophilus* Csm1 D575A D576A | 3892 | GGGAGAGGATCCATAAAGGAGG |
| | 3895 | ACCACCGGCATAGATAATGGA |
| | 3896 | TCCATTATCTATGCCGGTGGTGCAGCAGTTTTTGCAATTGGTAGCTGGC |
| | 3897 | CGCAGCCATATGTTAATCTTTACG |

**Table S4. Northern analysis oligonucleotide probes**

| Probe | ID | Oligonucleotide sequence 5'-3' |
|---|---|---|
| 5S rRNA | 4636 | TTCTGAATTCGGCATGGGGTCAGGTGG |
| C crRNA | 3206 | ATTTCTGAAGTGCTCTCAGCCGCAAGGACCGCATACTACAA |
| lpp 3' (Fig 1-2) | 3233 | GAACGTCGGAACGCATTGCGTTCACGTCGTTGCTCAGCT |
| lpp crRNA | 3026 | ATTTAGCTACTAAACTGGTACTGGGCGCGGTAATCCTGG |
| lpp 3' (Fig S1) | 3098 | ACAGCTGATCGATTTTAGCGTTGCTGGAGCAACCT |
| lpp 5' | 4673 | ATACCCTCTAGATTGAGTTAATCTCCATGTAGCG |
| lpp 3' (Fig 3) | 4750 | TGCGCCATTTTTCACTTCACAGGTACTATTACTT |
| cspE 3' | 4681 | GACGTATCTTACAGAGCGAT |
| ompF 5' | 3203 | CTGCCAGAATATTGCGCTTCATCATTATTTATTAC |
| rnpB 5' | 5236 | CGGCGACTGTCTGGTCAGCTTC |
| valV-valW | 3190 | CGGACGCAGGATGGTGCGTT |
